# The nanoscale organization of reticulon 4 shapes local endoplasmic reticulum structure *in situ*

**DOI:** 10.1101/2023.01.26.525608

**Authors:** Lukas A. Fuentes, Zach Marin, Jonathan Tyson, David Baddeley, Joerg Bewersdorf

**Author notes:** L. A. Fuentes and Z. Marin contributed equally to this paper.

## Abstract

The endoplasmic reticulum’s (ER) structure is directly linked to the many functions of the ER but its formation is not fully understood. We investigate how the ER-membrane curving protein reticulon 4 (Rtn4) localizes to and organizes in the membrane and how that affects local ER structure. We show a strong correlation between the local Rtn4 density and the local ER membrane curvature. Our data further reveal that the typical ER tubule possesses an elliptical cross-section with Rtn4 enriched at either end of the major axis. Rtn4 oligomers are linear-shaped, contain about five copies of the protein, and preferentially orient parallel to the tubule axis. Our observations support a mechanism in which oligomerization leads to an increase of the local Rtn4 concentration with each molecule increasing membrane curvature through a hairpin wedging mechanism. This quantitative analysis of Rtn4 and its effects on the ER membrane result in a new model of tubule shape as it relates to Rtn4.

**Summary:** Rtn4 forms linear-shaped oligomers that contain an average of five Rtn4 proteins, localize to the sides of elliptical tubules, prefer orientations near parallel to the tubule axis, and increase local curvature of the ER membrane by increasing local Rtn4 density.

## Introduction

The endoplasmic reticulum (ER) is the site of several critical cellular functions which are associated with specific ER substructures. These include the morphological regulation of other organelles by tubules (Abrisch et al., 2020; Friedman et al., 2011; Hoyer et al., 2018; Zheng et al., 2021), protein translation on rough ER sheets (Jan et al., 2014; Shibata et al., 2010; West et al., 2011), and lipid biogenesis in the tubule-rich ER-Golgi intermediate compartment (Glick and Nakano, 2009). The structure of the ER adapts to the needs of different cell types, displaying expansive sheets for enhanced protein expression and secretion or highly tubular ER in the case of androgen production (Lee et al., 2005; Zirkin and Papadopoulos, 2018). This critical link between ER structure and function has been made clear as disrupting the specialized processes of the ER in these cells consistently leads to perturbed ER structure (Lee et al., 2005; Zirkin and Papadopoulos, 2018). Recently, ER structure has been strongly linked to markers of disease state for obesity and diabetes in liver tissues by the remarkable discovery that these markers can be rescued to levels of non-diseased tissues by manipulation of ER structure alone (Parlakgül et al., 2022).

Investigations of ER morphology are, however, complicated by the fact that the sizes of typical ER features are below the diffraction limit and hence impossible to resolve by standard light microscopes. Studies making use of super-resolution microscopy (Nixon-Abell et al., 2016; Schroeder et al., 2019; Wang et al., 2022) or electron microscopy (Puhka et al., 2012; Terasaki et al., 2013; Zamponi et al., 2022) methods have highlighted the fact that ER structure is more complex than once thought, as structural details of the ER were not fully appreciated when visualized with conventional light microscopy methods. What were once thought to be continuous ER sheets have been revealed to often be intricate tubular matrices (Nixon-Abell et al., 2016) and sheets containing dynamic nanoholes (Schroeder et al., 2019). Additionally, two types of tubules have been described with the larger ones possessing attributes consistent with elliptical cross-sections (Wang et al., 2022). Another study using Airyscan microscopy and cryo-electron tomography revealed another sub-diffraction structure, termed ER ladders, found in developing axons (Zamponi et al., 2022).

Several proteins in the ER are responsible for promoting specific ER substructures. Sheet thickness is maintained by CLIMP63, while p-180 and kinectin have been speculated to maintain sheet flatness (Klopfenstein et al., 2001; Shen et al., 2019; Shibata et al., 2010). The reticulon (Rtn) and REEPs/DP1/Yop1p protein families are known to be responsible for stabilizing curvature in the ER membrane (Voeltz et al., 2006; Hu et al., 2008). Both protein families possess reticulon homology domains (RHD) with hairpin topologies that do not completely pass through the membrane bilayer (Voeltz et al., 2006; Zurek et al., 2011). This results in the proteins forming wedges in the ER membrane that displace more phospholipids in the cytosolic leaflet than the luminal leaflet. This wedging mechanism is likely a major contributor to the membrane curving function of these proteins. REEPs/DP1/Yop1p have been shown to form dimers that may organize into a splayed conformation to promote membrane curvature (Wang et al., 2021). Due to reticulons sharing a similar topology to REEPs, this mechanism has been considered for the reticulon proteins as well (Wang et al., 2021). However, reticulons and REEPs/DP1/Yop1p have also been shown to form higher-order homo-oligomers *in vitro*, and it was proposed that these oligomers form arches in the membrane to scaffold it into a curved topology (Shibata et al., 2008; Hu et al., 2008). However, due to the difficulty of visualizing such structures at the size scale of tens of nanometers, neither model has been confirmed yet by direct visualization of these complexes.

Reticulon 4 (Rtn4) is one of the best-studied proteins among the reticulon and REEPs/DP1/Yop1p families of proteins, especially in terms of its localization. In addition to its well-known partitioning to tubules and sheet edges (Shibata et al., 2010; Voeltz et al., 2006), Rtn4 generally localizes to areas of higher membrane curvature (Kiseleva et al., 2007) and nanoholes within ER sheets (Schroeder et al., 2019). In wider tubules it also displays two parallel lines along their sides (Wang et al., 2022). Rtn4’s several isoforms all possess a C-terminal RHD (Oertle and Schwab, 2003; Yang and Strittmatter, 2007; Schroeder et al., 2019) that is important for localization as well as Rtn4’s ability to curve membranes, which suggests all RHD-containing proteins share a general mechanism for shaping membranes (Zurek et al., 2011), though the exact mechanism remains unclear.

Despite the critical importance of ER structure on cellular function and health, there is still much we do not understand about how ER-shaping proteins form and maintain that structure. Additionally, Rtn4 has been implicated in neurological diseases such as Alzheimer’s disease, amyotrophic lateral sclerosis, and multiple sclerosis (He et al., 2004; Jokic et al., 2005; Karnezis et al., 2004; Chiurchiù et al., 2014). Gaining insight into Rtn4’s function and its effects on ER structure could help elucidate its role in these diseases.

In this study, we set out to understand how Rtn4 proteins are organized at the nanoscale *in situ* to perform their membrane-curving function and how that organization affects local ER structure. We make use of a suite of recently developed software tools (Barentine et al., 2018; Thevathasan et al., 2019; Marin et al., 2021) (Marin et al., in preparation) and super-resolution methods (Zhang et al., 2020; Chung et al., 2022) to accomplish this. Our results lead us to propose a refined model for Rtn4 localization, organization, and its effect on tubule shape.

## Results

### Rtn4 density determines local ER membrane curvature

We first set out to understand how Rtn4 distribution and local abundance can affect ER structure. Using Stimulated Emission Depletion (STED) microscopy, we imaged Rtn4, endogenously tagged with HaloTag at the C-termini of all isoforms by CRISPR Cas9 gene editing, in live U-2 OS cells **(Fig. S1, A-C)**. We additionally overexpressed Sec61β, an ER membrane protein, tagged with SNAP-tag to provide information about the underlying ER structure. Analysis of the difference in Rtn4 pixel intensities along tubules revealed that Rtn4 is significantly more abundant in tubules at the outer periphery of the cell than in those that are closer to the nucleus **(Fig. 1, A and B)**. We asked if this discrepancy in the abundance of Rtn4 in different regions of the ER is correlated with differences in the underlying structures. Specifically, we hypothesized that more Rtn4 present in a tubule leads to a smaller tubule diameter. From our live-cell STED images, we measured the diameters of tubules located at the periphery of the ER and compared them to the diameters of perinuclear tubules using our previously developed NEP-Fitting tool (Barentine et al., 2018). The peripheral (Rtn4-enriched) tubules displayed significantly smaller diameters on average **(Fig. 1, C and D)**, consistent with our hypothesis. However, to ensure that this difference in tubule diameters was due to varying amounts of Rtn4 alone, we imaged overexpressed Sec61β tagged with HaloTag in both live U-2 OS and live U-2 OS Rtn4 knock-out cells (Schroeder et al., 2019) and compared the diameters of their ER tubules **(Fig. 1, E-G)**. Our analysis showed a significant difference between them with U-2 OS and U-2 OS Rtn4 knock-out cells possessing 94 nm and 119 nm diameter tubules on average, respectively. Interestingly, our measured tubule diameters showed a broad distribution with 90% of diameters covering a range of ∼50 to ∼130 nm, which is consistent with our previous analyses (Barentine et al., 2018; Schroeder et al., 2019) but not the recent observation of two distinct ER tubule diameters (Wang et al., 2022). Altogether, these results strongly suggest that higher Rtn4 abundance leads to thinner tubule diameters, consistent with earlier findings of *in vitro* studies (Hu et al., 2008).

**Fig. 1.**
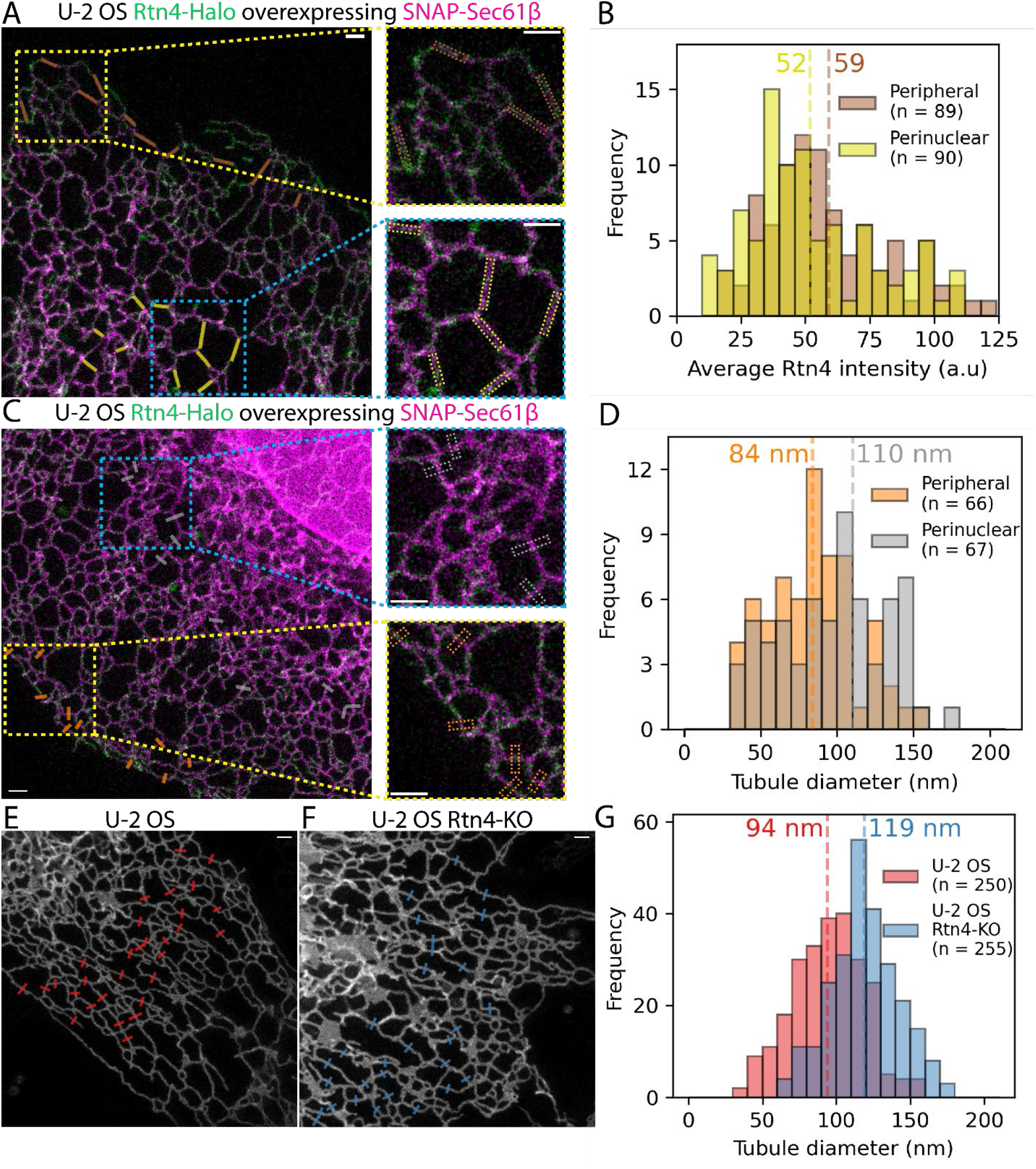
Effects of Rtn4’s distribution on microscale ER structure. (A, C) Two-color Live-Cell STED images of U-2 OS Rtn4-Halo (green) CRISPR cells overexpressing SNAP-Sec61β (magenta). (A) Brown and yellow boxes are highlighting where regions of interest were made for Rtn4 pixel intensity measurements in peripheral and perinuclear tubules, respectively. (B) Histogram of average Rtn4 pixel intensity measured along tubules (p < 0.05; Wilcoxon rank-sum statistic). (C) Orange and grey boxes are highlighting where line plots were made for tubule diameter measurements using the NEP fitting method based on SNAP-Sec61β fluorescence for peripheral and perinuclear tubules, respectively. (D) Histogram of diameters of peripheral and perinuclear tubules (p < 0.01; Wilcoxon rank-sum statistic). (E, F) STED images of Halo-Sec61β overexpressed in U-2 OS and U-2 OS Rtn4-KO cells, respectively. Red and blue boxes represent where line plots were created for tubule diameter measurements for each dataset. (G) Diameters of U-2 OS and U-2 OS Rtn4-KO cells’ tubules (p < 0.001; Wilcoxon rank-sum statistic). (B, D, G) Hashed vertical lines represent the average values which appear near each line. (A, C, E, F) The scale bars represent 1 μm.

To better understand how the density of Rtn4 affects ER nanostructure *locally*, we imaged endogenous Rtn4 and overexpressed mCherry-Sec61β in U-2 OS cells using 3D single-molecule localization microscopy (3D-SMLM) on our custom-built 4Pi-SMS microscope that enables isotropic ∼20-nm resolution in multiple colors (Huang et al., 2016; Zhang et al., 2020). Employing a software tool that generates accurate 3D surfaces from point-cloud data (Marin *et al*., in preparation), we created 3D surfaces based on the mCherry-Sec61β point-cloud to serve as a proxy for the ER membrane **(Fig. 2 A; Video 1)**. We calculate the mean surface curvature **(Fig. 2 B)** as well as the Rtn4 density within 50 nm of each vertex on the surface **(Fig. 2 C; Fig. S2)**. Locations of high Rtn4 density consistently correlate with locations of high mean curvature **(Fig. 2, B and C; blue arrowheads)**. To support this observation quantitatively, we examined Rtn4 density as a function of the curvature of its tubular cross section (the maximal principal curvature at each vertex on a tubule). We normalized the Rtn4 density by the frequency of the maximal principal curvatures that each nearby surface vertex possessed. This analysis revealed that as the maximal principal curvature increases, so does the Rtn4 density **(Fig. 2 D)**. The Rtn4 density peaks at a curvature of 0.073 nm^-1^, which translates to a tubule diameter of ∼27 nm (equal to two times the radius of 1/0.073 nm). This tubule size is consistent with the smallest reported ER tubules (Terasaki, 2018; Zamponi et al., 2022) as well as theoretical models of ER sheet edge radii of curvature (Shibata et al., 2010). These results suggest that the local density of Rtn4 has a significant role in determining the local membrane curvature.

**Fig. 2.**
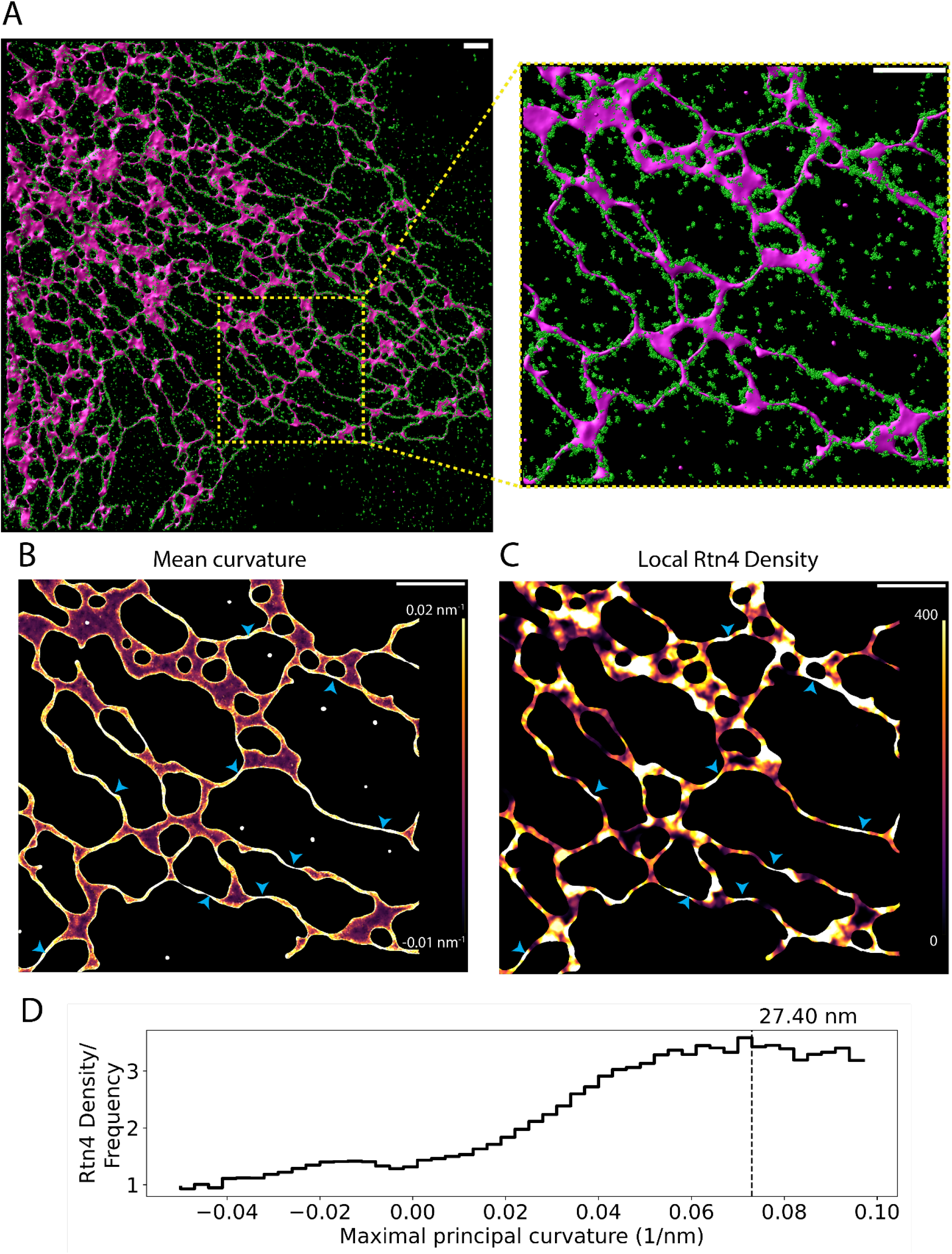
Effects of Rtn4’s distribution on nanoscale ER structure. (A) 3D data of Rtn4 localizations (green spheres) and a 3D surface based on Sec61β localizations (magenta-shaded surface). (B) 3D surface colored by mean curvature which ranges from -0.01 to 0.02 nm^-1^. (C) 3D surface colored by the local Rtn4 density within 50 nm of each vertex on the 3D surface ranging from 0 to 400 localizations. The blue arrowheads highlight locations on the surface that possess a high Rtn4 density nearby. (D) Histogram of the frequency of Rtn4 localizations occurring within 50 nm of a given vertex after normalizing for the frequency of the vertices maximal principal curvatures (the curvature of the circular cross-section of the tubule). The hashed line represents the curvature where the Rtn4 density is the highest. The value above the hashed line represents the diameter of a hypothetical tubule with such a curvature. The scale bars represent 1 μm.

### ER tubules have elliptical cross-sections and Rtn4 localizes to their sides

As expected, we observed from our 3D data that Rtn4 preferentially localizes to the edges of sheet-like structures, but not their flat top and bottom surfaces. For tubules, Rtn4 also appears to prefer their sides, in the x-y plane of the sample, over the tops and bottoms of them **(Fig. 2 A)**, confirming a recently reported observation by Wang et al. (Wang et al., 2022). Most ER models depict tubules with circular cross-sections (Goyal and Blackstone, 2013; Georgiades et al., 2017; Shibata et al., 2010; Wang and Rapoport, 2019; Chen et al., 2013; Lin et al., 2012; Shemesh et al., 2014; Obara et al., 2022; Hu et al., 2008), and one would expect a random distribution of Rtn4 around such tubules. However, as recently suggested (Wang et al., 2022), tubules with elliptical cross-sections, where Rtn4 preferentially locates to the regions of higher membrane curvature, could explain our observation. To investigate this in more detail, we considered four possible models **(Fig. 3 A)**: (i) tubules with circular cross-section and random Rtn4 distribution; (ii) tubules with circular cross-section and enrichment of Rtn4 at the tubule sides; (iii) tubules with elliptical cross-sections, with the long axis in the x-y plane, and random Rtn4 distribution; and (iv) tubules with elliptical cross-sections and Rtn4 enriched at the tubule sides. To test which of these models is consistent with our data, we performed 3D-SMLM using our 4Pi-SMS instrument (Zhang et al., 2020; Huang et al., 2016) on three different samples. The first one was U-2 OS cells overexpressing mCherry-Sec61β labeled with an anti-RFP nanobody and the endogenous Rtn4 labeled with anti-Rtn4 primary, labeling the Rtn4B/D isoforms that are known to be expressed in U-2 OS cells (Schroeder et al., 2019), and secondary antibodies. The other two were U-2 OS cells overexpressing either mEmerald-Sec61β or GFP-KDEL. These proteins were double-labeled, first with an anti-GFP nanobody in one color and then with an anti-GFP primary and secondary antibodies in a second color, to mimic the labeling approach of the first sample. Data generated from localizing the nanobody fluorescence in each sample was used to create 3D surfaces which were then used to generate ‘skeletons’ of the ER network by applying mean curvature flow (Tagliasacchi et al., 2012) **(Fig. S3, A and B;** see methods). These skeletons provide critical information about the center and major axis of each tubule in the datasets **(Fig. 3, B and D)**. Using this information and the localizations from the secondary antibody fluorescence, we calculated the angles between the vectors pointing from each secondary-antibody localization to the closest point on the skeleton and the vector denoting the vertical (z) direction of the sample. We refer to these angles as the cross-sectional angles φ **(Fig. 3 A; model (i))**. We compared the distributions of φ in the three samples to that of simulated points randomly distributed on a circular ring around the skeletons **(Fig. 3, E-G; Fig. S3 C)**. We found that Rtn4, Sec61β, and KDEL are all enriched around ± 90° from the z-direction of the cross-sections when compared to simulated random distributions. Since KDEL is located in the ER lumen and no preference of Sec61β for specific membrane topology has been reported, we conclude from this observation that ER tubules are on average elliptical in shape, ruling out the first two models. Additionally, the three φ distributions are significantly different from each other with Rtn4 being the most enriched at the sides, followed by Sec61β and then KDEL **(Fig. S4 A)**. To show these results are not an artifact caused by the large size of primary and secondary antibodies, we repeated the φ analysis on the nanobody localizations of Sec61β and KDEL. These φ distributions show the same enrichment at the sides of tubules **(Fig. S4, B and C)**. The only model that is consistent with these observations is model (iv) **(Fig. 3 A)**, i.e., tubules with elliptical cross-sections and Rtn4 localized to the sides where the local curvature is highest. Indeed, visualizing cross-sections of tubules in our two-color 3D-SMLM data of Rtn4 and Sec61β revealed cross-sections with elliptical shapes and distributions of these two proteins consistent with model (iv) **(Fig. 3, H-N)**.

**Fig. 3.**
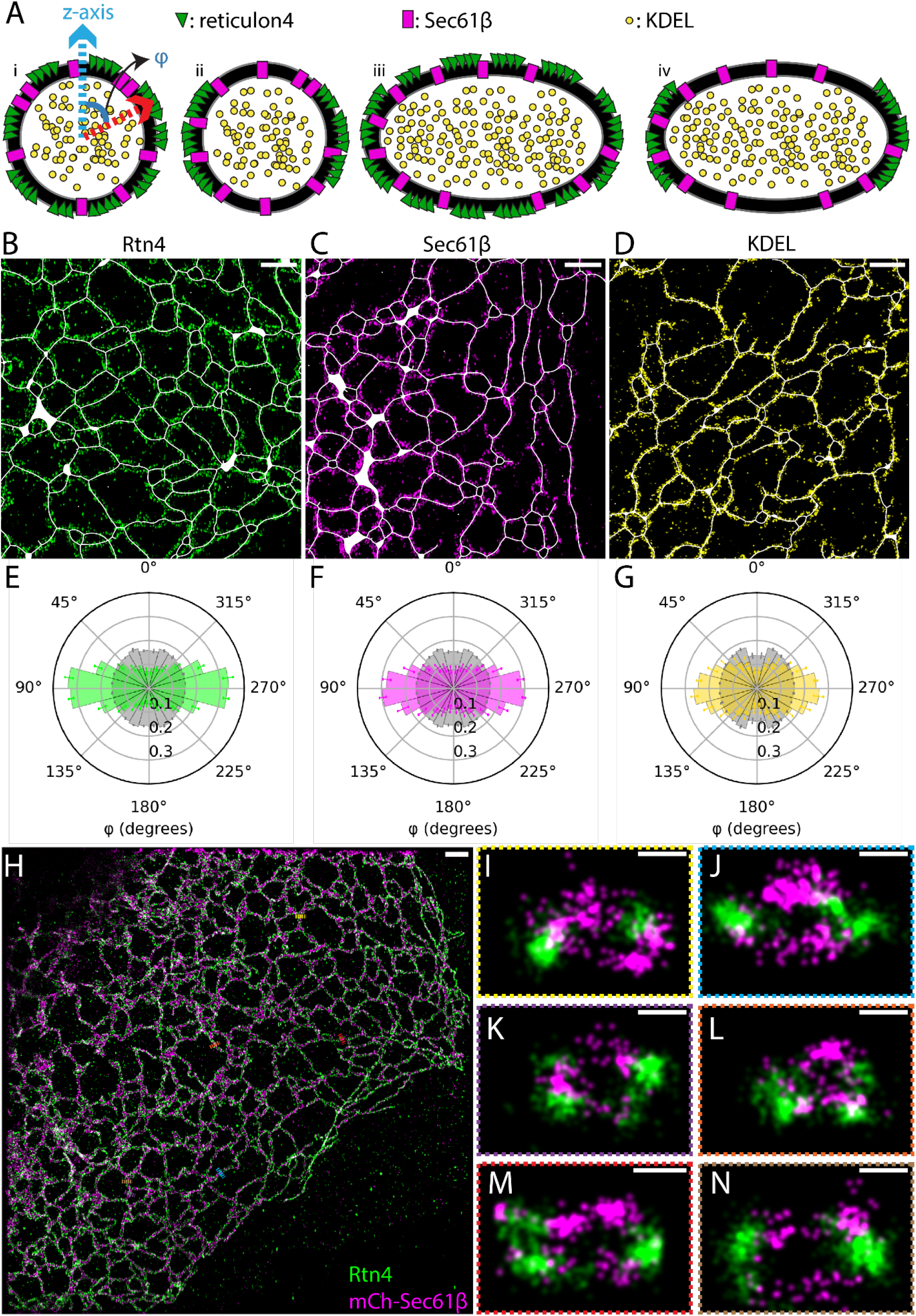
Angular distribution of ER proteins around tubule cross-sections. (A) Possible models of ER protein distributions in tubule cross-sections. Model (i) shows the vectors that angle φ is being calculated between. (B-D) Representative regions of interest taken from datasets of each protein. Skeletons, Rtn4, Sec61β, and KDEL are displayed in white, green, magenta, and yellow, respectively. (E-G) Polar histograms of φ. The Rtn4, Sec61β, and KDEL observed data is shown by the green, magenta, and yellow histograms, respectively. The grey histograms are from simulated datasets of points randomly distributed in a circular ring around the skeleton that was created based on each set of observed data (p <<< 0.001 for each protein’s distribution compared to their simulated random distributions; Wilcoxon rank-sum statistic). (H) Overview of a U-2 OS cell overexpressing mCherry-Sec61β (magenta) labeled with an anti-RFP nanobody and endogenous Rtn4 (green) labeled with primary and secondary antibodies. (I-N) Cross-sections of ER tubules with 3D views orthogonal to the tubule axis. Data was recorded with a 4Pi-SMS microscope. Scale bar: 1 μm (B-D and H); 50 nm (I-N).

### Rtn4 oligomers contain an average of five copies

A visual comparison of Sec61β and Rtn4 point clouds from our 3D SMLM data of ER tubules clearly shows Rtn4 being clustered while Sec61β appears diffusely localized **(Fig. S5)**, confirming earlier reports of Rtn4 oligomerization (Shibata et al., 2008). To gain insight into the organization of individual Rtn4 oligomers, we determined how many copies are in a typical oligomer using the powerful SMLM variant of fluorogenic DNA-PAINT (Chung et al., 2022) and single-molecule counting. Single-molecule counting results can be confounded by the occurrence of multiple events per protein and incomplete labeling (Baddeley and Bewersdorf, 2018). Following an established procedure (Thevathasan et al., 2019), we used the nucleoporin Nup96 as a standard to resolve these complications. We mixed a U-2 OS Nup96-mEGFP cell line (Thevathasan et al., 2019) with a U-2 OS Rtn4-mEGFP cell line **(Fig. S1, D-F)**, both endogenously tagged using CRISPR including all isoforms of Rtn4, into one sample and labeled the proteins of interest with an anti-GFP nanobody conjugated to a docking strand for fluorogenic DNA-PAINT probes. This allowed us to image both Nup96 and Rtn4 in one field of view, ensuring that all sample preparation and imaging conditions between both proteins were identical **(Fig. 4 A)**. The Nup96-mEGFP localizations were separated into their own regions of interest (ROIs) and fed into a set of software tools specifically designed for analyzing single-molecule localization data of Nup96, providing information about the effective labeling efficiency of our sample and the number of localizations per protein (Thevathasan et al., 2019) **(Fig. 4, B and C)**. We then created a ROIs of the Rtn4-mEGFP localizations and segmented them into clusters using density-based spatial clustering of applications with noise (DBSCAN) (Ester et al., 1996; Schubert et al., 2017) **(Fig. 4, D and E)**. Using the results of the Nup96-mEGFP counting analysis to calibrate the segmented clusters of Rtn4-mEGFP showed that a typical cluster, which we are considering to be equivalent to an oligomer, of Rtn4-mEGFP contains ∼5 copies of the protein (average of the median values across three experiments: 5.26) **(Fig. 4 F; red hashed line)**. This result is consistent with previously reported *in vitro* studies (Shibata et al., 2008), and confirms that Rtn4 also forms oligomers around this size *in situ*.

**Fig. 4.**
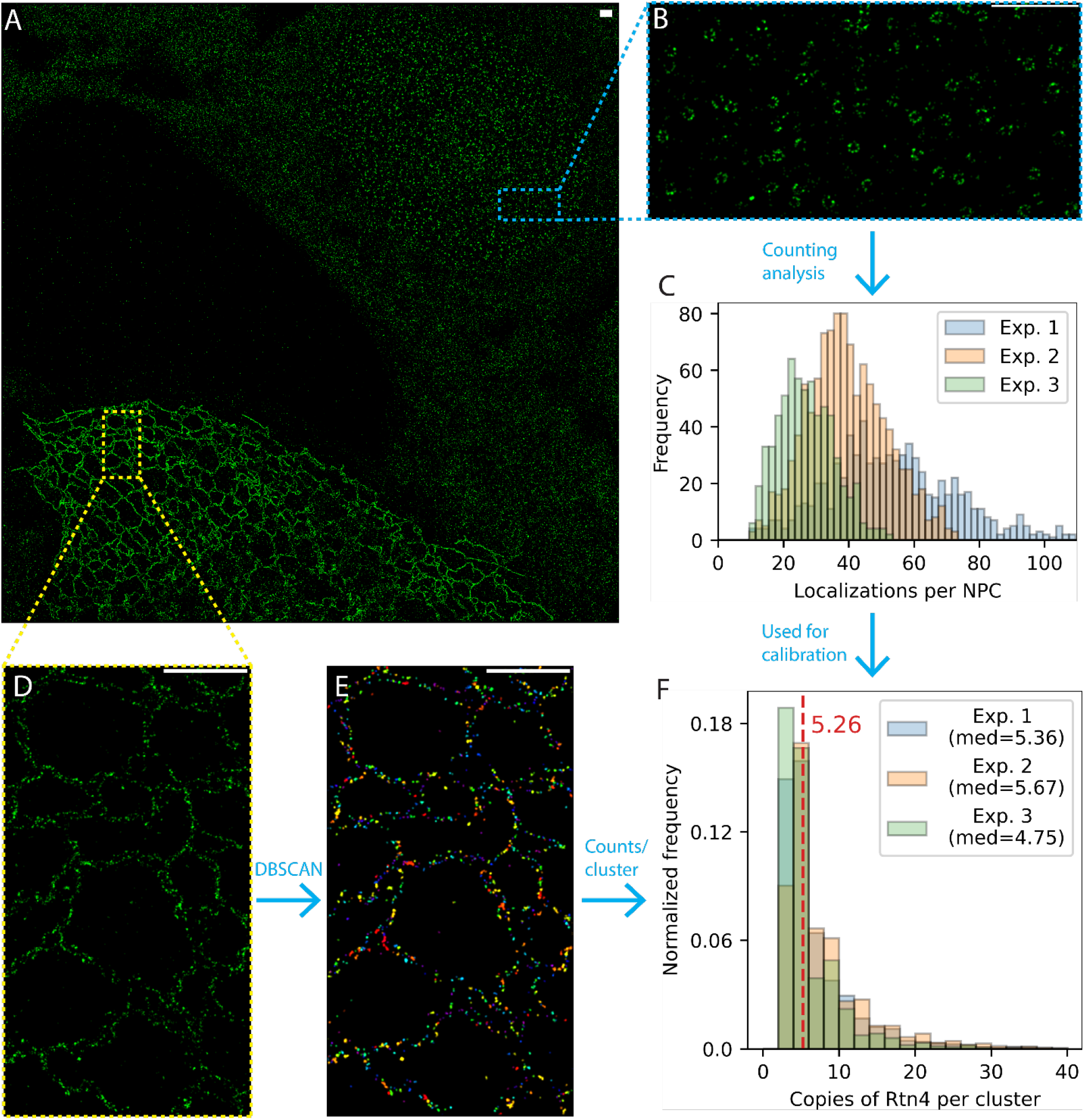
Counting the number of Rtn4 copies per oligomer. (A) Representative whole field-of-view of a dataset. A U-2 OS Rtn4-mEGFP cell and a U-2 OS Nup96-mEGFP cell are located in the bottom left and the upper right of the image, respectively. (B) Zoom-in of Nup96-mEGFP. (C) Histograms from each experiment of the number of localizations per nuclear pore complex. (D) Zoom-in of Rtn4-mEGFP. (E) Same zoom-in as (D) after segmenting clusters of Rtn4 localizations using DBSCAN. Clusters are colored uniquely. (F) Histograms from each experiment of the number of Rtn4 copies per cluster of Rtn4 localizations. Scale bars represent 1 μm.

### Rtn4 forms linear-shaped clusters enriched at orientations near parallel to the tubule axis

We next sought to understand how Rtn4 oligomers are organized in the ER membrane. We used DBSCAN on the 3D Rtn4 point clouds to segment the localizations into clusters **(Fig. 5 A; and Video 2)**. Analysis of φ for the centers of each cluster **(Fig. 5 B)** showed a similar result to the angular distribution of the Rtn4 localizations directly **(Fig. 3 E)**. Specifically, cluster centers displayed a strong enrichment at tubule sides. We next used principal component analysis (PCA) (Jollife and Cadima, 2016) to determine the principal axis of each cluster with the highest eigenvalue, which we refer to as the major axis. PCA also allowed us to analyze the shape of Rtn4 clusters by comparing the anisotropy of observed clusters to simulated clusters that were circular-or linear-shaped **(Fig. 5, C and D; Videos 3 and 4)**. Each simulated cluster mimicked an observed cluster by having the same center location, radius of gyration, and number of localizations, each of which had an identical localization precision to the real data incorporated in their simulated positions. The observed clusters had anisotropy distributions much more similar to the linear-shaped simulated clusters than the circular-shaped ones **(Fig. 5 E)**. This suggests that the Rtn4 oligomers are organized linearly with discernable major axes rather than forming circular oligomers that lack such axes. The clusters clearly did not consistently appear as arches oriented orthogonal to the tubule axis (Shibata et al., 2008; Hu et al., 2008) which led us to ask if the clusters’ major axes adopted consistent orientations relative to the tubule axes. To help answer this question, we simulated clusters with the same locations as the observed clusters, but with randomly oriented major axes, with the one restriction that these axes must be tangential to the ER surface **(Fig. 5 F; and Video 5)**. We further projected the vectors of the observed major axes of the clusters onto a plane approximately tangential to the 3D surface **(Fig. 5 G; red vector)** and calculated the angles ψ between them and vectors in the same plane but perpendicular to the tubule axis **(Fig. 5 G; blue vector)**. Comparing the distributions of ψ for the observed and simulated clusters shows that the observed clusters are strongly enriched at orientations close to parallel with the tubule axis (± 90°) and depleted at orientations close to orthogonal to the tubule axis **(Fig. 5H)**. This orientation preference of the observed clusters only disappears when limiting the analysis to the smallest clusters where the cluster orientations cannot be resolved by our microscope (**Fig. S6 A**).

**Fig. 5.**
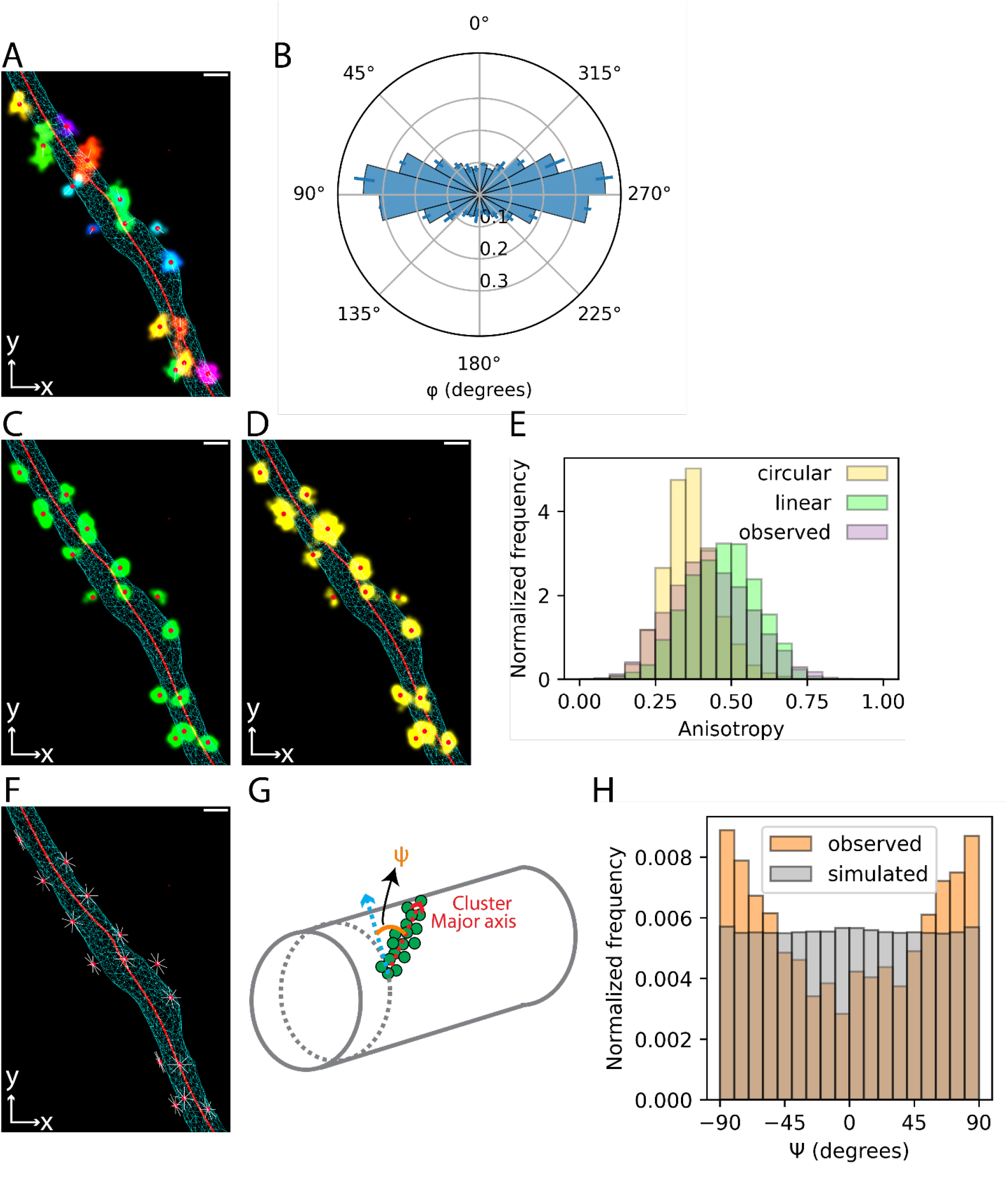
Position, shape, and orientation of Rtn4 oligomers. (A) An ER tubule showing the 3D surface as a cyan wireframe, the skeleton as a red line, each observed Rtn4 cluster colored uniquely, the centers of clusters as red circles, and the major axis of each cluster as a white line. (A) Polar histogram of φ for the centers of Rtn4 clusters. (C) Similar to (A) but with linear-shaped simulated clusters (green) instead of observed clusters. (D) Similar to (A) but with circular-shaped simulated clusters (yellow) instead of observed clusters. (E) Histograms of the anisotropy of observed clusters (lavender), linear-shaped simulated clusters (green) and circular-shaped simulated clusters (yellow). (F) Similar to (A) excluding observed clusters and with simulated cluster major axes shown that are randomly oriented and tangential to the tubule surface. (G) Model of the vectors that ψ is being calculated between. ψ is calculated between the major axes of the clusters (red arrow) and vectors orthogonal to the tubule axis (blue arrow). See methods for more details. (H) Histogram of ψ for observed clusters and clusters simulated to possess all orientations tangential to the tubule surface (F).

## Discussion

Using super-resolution imaging and novel software tools, we were able to elucidate the organization of Rtn4 at the nanoscale in U-2 OS cells and how that affects local ER structure. Live-cell STED microscopy data and mean curvature analysis of 3D surfaces generated from 3D point-cloud data showed that the local curvature of the ER membrane in cells increases with the local density of Rtn4. Our analyses of tubule cross-sections revealed that most ER tubules are elliptical with Rtn4 localized specifically to their sides where the curvature is greatest. Oligomers of Rtn4 in cells were found to contain ∼5 copies of the protein based on single-molecule counting data. Lastly, we analyzed the shape and orientation of Rtn4 oligomers on ER tubules and could show that they are linear-shaped and prefer orientations parallel to the tubule axis. These findings result in a refined model of Rtn4 organization, localization, and the resulting shape of ER tubules **(Fig. 6)**.

**Fig. 6.**
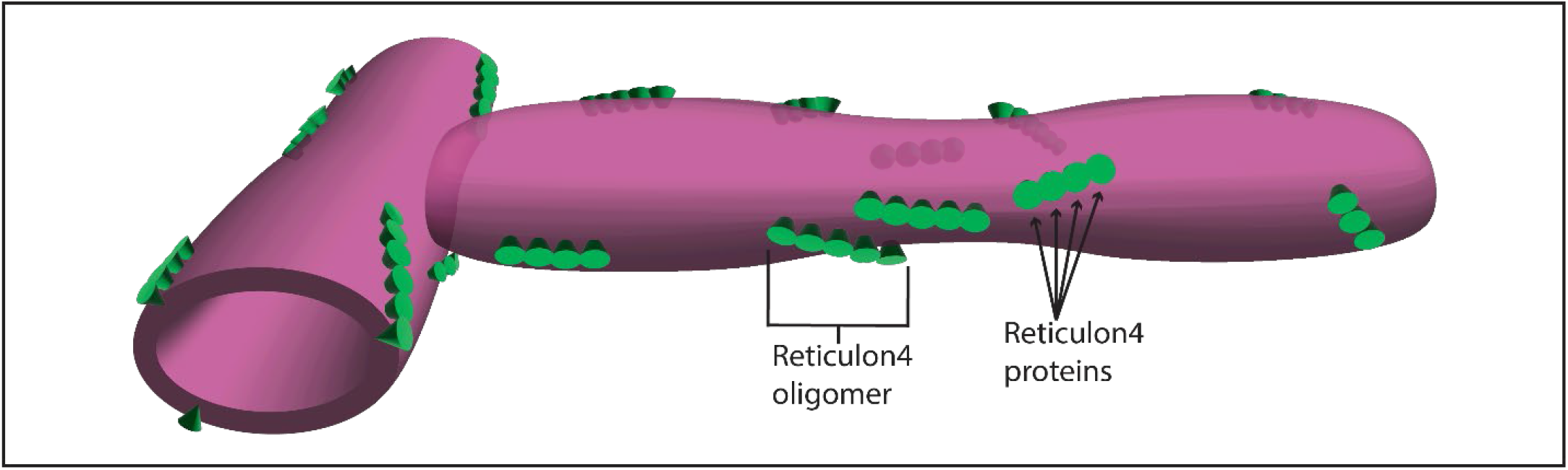
Model of Rtn4 organization and its effects on tubule structure. Individual Rtn4 proteins are depicted as green cones on a magenta tubule surface.

Until recently, tubule models have all assumed circular cross-sections (Goyal and Blackstone, 2013; Georgiades et al., 2017; Shibata et al., 2010; Wang and Rapoport, 2019; Chen et al., 2013; Lin et al., 2012; Shemesh et al., 2014; Hu et al., 2008; Obara et al., 2022). These were likely heavily influenced by electron microscopy data as well as *in vitro* reconstitutions of ER networks which displayed very narrow, and likely round in cross-section, tubules due to the presence of membrane curving proteins at concentrations ∼20x higher than physiological concentrations (Hu et al., 2008). Our data show that the tubule diameter decreases as the local concentration of Rtn4 increases. Thus, it is easy to see why *in vitro* reconstitutions of the ER using high concentrations of membrane curving proteins result in tubules with circular cross-sections. It is likely that some tubules in cells possess enough Rtn4 to take on this shape, as has also been suggested by others (Wang et al., 2022). According to our results, such tubules would most likely occur in the periphery of the ER. However, the vast majority of tubules we observe in U-2 OS cells appear to have an elliptical shape, and surveying past EM studies of ER structure has revealed datasets with elliptical tubules as well (Puhka et al., 2012; Nixon-Abell et al., 2016).

With elliptical tubules, the line between what is a tubule and what is a sheet becomes blurred. Recent studies of ER structure have made it clear that this differentiation is not as binary as once thought. Instead, the ER displays a spectrum of morphologies (Nixon-Abell et al., 2016; Schroeder et al., 2019; Puhka et al., 2012). Thus, it may be more appropriate to think of the ER as being composed of a spectrum of sheets including continuous sheets, fenestrated sheets, and now (structurally speaking) very narrow sheets, referred to as ribbon-like ER sheets by Wang et al. (Wang et al., 2022), that have traditionally been thought of as tubules.

We consistently observed the major axes of elliptical tubule cross-sections lying in the x-y plane. We do not believe this to be universally true, but rather a side-effect of focusing our imaging efforts in the lamellipodia of very flat cells. It is unclear what causes this phenomenon. One possibility is that the plasma membrane is physically ‘squeezing’ the peripheral ER, thus restricting the orientations of the elliptical cross-sections. Considering this, it is important to note that our diameter measurements on STED data are measuring the width of tubules in the x-y plane which, in our case, is the major axis of the elliptical cross-sections.

It was recently suggested that ER tubules exist in two distinct forms with consistent diameters of ∼105 nm and ∼50 nm, with U-2 OS cells’ tubules existing predominantly in the ∼105 nm form (Wang et al., 2022). However, our diameter measurements here and in previous analyses (Barentine et al., 2018; Schroeder et al., 2019) do not show consistent narrowly distributed tubule measurements with peaks around these values. Instead, they show broad ranges that include tubules with ∼105 nm and ∼50 nm amongst many other sizes.

We have shown Rtn4’s strong enrichment at the sides of tubules where the curvature is highest in the elliptical cross-sections. This was observed for both individual localizations of Rtn4 and the centers of Rtn4 clusters which is indicative of Rtn4 not forming oligomers of different sizes at different positions on the tubule. Plotting the measured positional angles φ of the clusters against their orientational angles ψ, we also see that the orientation of the clusters does not depend on where the clusters are located on the tubules **(Fig. S6 B)**.

We observe regions of high membrane curvature that exhibit low local Rtn4 density **(Fig. 2, B and C)**. This can be explained by the presence of other membrane curving proteins within the ER (Kumar et al., 2021; Voeltz et al., 2006; Shibata et al., 2008, 2010; Park et al., 2010; Mochida et al., 2015; Dong et al., 2018) as well as by potential external forces and thermodynamic fluctuations of the membrane. In essence, a high concentration of Rtn4 will lead to high membrane curvature, but high membrane curvature is not always caused by high Rtn4 concentration.

We have also shown that Rtn4 organizes into oligomers of around five copies of the protein. However, it is not strictly a pentamer. As our single-molecule counting results show, there is a distribution of sizes. To get a sense of how monomers of Rtn4 are organized within an oligomer, we simulated point clouds of Rtn4 oligomers possessing specific distances between monomers positioned in a line (see methods) and compared their anisotropies to those of observed Rtn4 oligomers **(Fig. S7)**. We calculated the theoretical diameter of a Rtn4b molecules to be ∼4.5 nm, assuming they are roughly spherical. Using this value as the inter-monomer distance in the simulation resulted in remarkably similar distributions between the observed and simulated clusters’ anisotropies **(Fig. S7 C)**, while simulations for half or double that distance showed strong discrepancies **(Fig. S7, A and D)**. Decreasing the inter-monomer distance to 3.75 nm resulted in nearly perfect overlap between simulation and experimental results **(Fig. S7 B)**. This simple simulation supports the idea that Rtn4 oligomers are composed of monomers that roughly organize in a linear fashion.

Until now, it was attractive to think that Rtn4 oligomers formed arches around tubules and sheet edges to scaffold the membrane into a curvature (Shibata et al., 2008; Hu et al., 2008). However, we have seen the majority of oligomers at orientations near parallel to tubule axes, which is the opposite of what one would expect for arching oligomer scaffolds. Our results are also inconsistent with the possibility of Rtn4 forming predominantly dimers that could adopt a splayed confirmation (Wang et al., 2021). We therefore believe that the oligomerization of Rtn4 in and of itself does not contribute to membrane curvature and instead serves as a mechanism to increase local Rtn4 concentration, which our results show correlates with higher curvature.

This further supports a model in which the membrane curvature is primarily induced by a hairpin wedging mechanism (Zurek et al., 2011; Voeltz et al., 2006). This explains both the importance of Rtn4’s oligomerization and the lack of its oligomers orienting in a way that would be amenable to a scaffolding mechanism. Consistent with this idea, long transmembrane domain mutants of Rtn4’s RHD have been reported to still form oligomers while lacking the ability to curve ER membrane (Zurek et al., 2011). We assume that the membrane curvature generated by Rtn4 monomers is in a single direction and that they are oriented in a specific direction within oligomers. Considering that oligomers preferentially orient parallel to the tubule axis, this model suggests that Rtn4 monomers generate curvature in the direction perpendicular to the major axis of oligomers.

Given Rtn4’s shared characteristics with other known ER membrane-curving proteins (Zurek et al., 2011; Voeltz et al., 2006; Shibata et al., 2008; Hu et al., 2008), it is reasonable to think that this model could be generalized to other reticulons and the REEPs/DP1/Yop1p family of proteins. Additionally, considering the many links between ER structure and disease (Sharoar et al., 2016; Parlakgül et al., 2022), some implicating Rtn4 directly (Zheng et al., 2022; Zhang et al., 2021; Kulczyńska-Przybik et al., 2021b; Boghdadi et al., 2018; Kulczyńska-Przybik et al., 2021a; Chiurchiù et al., 2014), the results of this study may provide additional insight in this regard.

## Materials and methods

### Cell culture

U-2 OS (ATCC; HTB-96; Lot # 70008732) were cultured in McCoy’s 5A (Gibco; 16600-082) + 10% FBS (Gibco; 10438-026). U-2 OS Rtn4-mEGFP, U-2 OS Rtn4-Halo, and the U-2 OS Nup96-mEGFP (Cell Lines Service; product number: 300174) CRISPR cell lines were cultured in the same way. Subculturing was performed using 0.05% Trypsin (Gibco; 25200056). Transfections were performed using a Super Electroporator NEPA21 Type II (Nepa Gene). Cells were concentrated to 1 million cells in 90 μL in an electroporation cuvette (Bulldog Bio; 12358-346) to which 10 μL of ∼1 μg/μL of plasmid DNA was added. Cells were electroporated with settings described previously (Schroeder et al., 2019). Prior to seeding cells, the coverslips for fixed samples were treated with ozone for 30 minutes, the ibidi slides for DNA PAINT were cleaned with plasma for 5 minutes (Harrick Plasma; PDC-001), and the dishes for live cells were cleaned by placing them in KOH and sonicating them in a water bath for 15 minutes. Cells were then seeded onto 30-mm round coverslips (Bioptechs; 30-1313-03192) for fixed 4Pi-SMS samples, ibidi slides (ibidi; 80607) for fluorogenic DNA PAINT single-molecule counting, or MatTek dishes for live STED samples (MatTek; P35G-1.5-20-C). Cells were allowed to adhere to the coverglass and reach desirable transient expression levels for 16-24 hours post-transfection before proceeding with sample preparation.

### Plasmids

GFP-KDEL was cloned as previously described (Merta et al., 2021). SNAP-Sec61β and Halo-Sec61β were cloned as previously described (Bottanelli et al., 2016). mEmerald-Sec61β and mCherry-Sec61β were acquired from Addgene (plasmids 54249 and 49155, respectively).

### CRISPR gene editing

The U-2 OS Rtn4-Halo cell line was developed using Cas9 gene editing. A gRNA described previously to generate a U-2 OS Rnt4-SNAP cell line was used in the generation of this cell line as well (5’-AAACGCCCAAAATAATTAGTAGG-3′; the PAM site is underlined) (Schroeder et al., 2019). The homology-dependent repair (HDR) template from that study was modified to replace the SNAP-tag sequence with the HaloTag sequence. This was accomplished using a 2-fragment Gibson Assembly using an NEBuilder reaction (New England Biolabs; E5520S). The fragments were amplified using PCR with the following primers (uppercase indicates Gibson overlapping bases):

Halo-Forward: 5’-GTCGCCACCATGGCAGAAATcggtactgg-3’

Halo-Reverse: 5’-GCTTTAGCCGGAAATCTCGagcgtc-3’

Vector-Forward: 5’-CGAGATTTCCGGCTAAAGCggccgcgactctagatc-3’

Vector-Reverse: 5’-ATTTCTGCCATGGTGGCGACcggtggatc-3’

U-2 OS (ATCC; HTB-96; Lot # 70008732) were transfected with the gRNA, HaloTag HDR template, and pSpCas9 (Addgene; 48137) via Lipofectamine 2000 in the well of a 6-well plate. Transfected cells were expanded for 1 week before selection with G418 began. After 9 days of selection, cells were labeled with SiR-chloroalkane for 1 hour followed by 3x washes with warm media and a 1-hour recovery at 37 °C. The cells were immediately sorted using FACS to obtain monoclonal cell lines. Cells were first screened visually for fluorescence consistent with Rtn4 localization. Then the success and zygosity of gene-editing was assessed by western blotting.

The U-2 OS Rtn4-mEGFP cell line was developed using Cas9 nickase D10A, to increase the chances of developing a homozygously tagged cell line (Koch et al., 2018). Following the protocol from Koch *et al*., a plasmid was cloned via T4 ligase using annealed oligos for the paired gRNA sequences and pX335-U6-Chimeric_BB-CBh-hSpCas9n(D10A) plasmid (Addgene; 42335) cut with BbsI restriction enzyme. The gRNA sequences were as follows: sense: 5’-AAACGCCCAAAATAATTAGT-3’; anti-sense: 5’-CAGCTTTGCGCTTCAATCCA-3’. A HDR template vector containing 800 bp homology arms flanking the RTN4 locus (Gene ID: 57142) was ordered for Genewiz. The mEGFP tag was cloned into the vector using a 2-fragment Gibson assembly with the following primers (uppercase indicates Gibson overlapping bases):

mEGFP-Forward: 5’-GGTGAGCAAGGGCGAGGAGCtgttcac-3’

mEGFP-Reverse: 5’-GGCGTTTTCACTTGTACAGCtcgtccatgc-3’

Donor-Vector-Forward: 5’-GCTGTACAAGTGAAAACGCCcaaaataattagtag-3’ Donor-Vector-Reverse: 5’-GCTCCTCGCCCTTGCTCACCatggtgg-3’

U-2 OS (ATCC; HTB-96; Lot # 70008732) were transfected with the pX335-U6-Chimeric-sense-gRNA, pX335-U6-Chimeric-antisense-gRNA, and mEGFP HDR template plasmids using JetPrime (Polyplus transfection; 114-07). Cells were expanded for 6 days before sorting with FACS to obtain monoclonal cell lines. Cells were expanded post-FACS for 3 weeks before visually screening for fluorescence consistent with Rtn4 localization. The success and zygosity of gene-editing were assessed by western blotting.

### Immunofluorescence

U-2 OS cells (ATCC; HTB-96; Lot # 70008732), or cell lines generated from this cell line, other than U-2 OS Nup96-mEGFP, were used for every experiment. All incubations for immunofluorescence samples were done at room temperature with the sample rocking, unless otherwise noted. Cells were fixed in 3% paraformaldehyde + 0.1% glutaraldehyde in 1xPBS for 15 minutes. The fixation was then quenched with 0.1% sodium borohydride in 1xPBS for 7 minutes followed by 100 mM glycine in 1xPBS for 10 minutes. Samples were rinsed three times with 1xPBS and permeabilized with room temperature permeabilization buffer (0.3% IGEPAL CA-630 [Sigma-Aldrich; I8896], 0.05% Triton X-100 [Sigma-Aldrich; T8787], 0.1% BSA [Jackson ImmunoResearch; 001-000-162] in 1xPBS) for 3 minutes followed by another three washes with 1xPBS. Samples were blocked for 1 hour with block buffer (0.05% IGEPAL CA-630, 0.05% Triton X-100, 5% goat normal serum [Jackson ImmunoResearch; 005-000-121] in 1xPBS). Samples were incubated with primaries/nanobodies diluted in block buffer for 16 hours at 4 °C on a rocker. Excess primaries/nanobodies were washed away with three 5-minute incubations with wash buffer (0.05% IGEPAL CA-630, 0.05% Triton X-100, 0.2% BSA in 1xPBS) before incubating samples with secondaries diluted in block buffer for 1 hour. Excess secondaries were washed away with three 5-minute incubations with wash buffer. Finally, samples were rinsed three times with 1xPBS. 4Pi-SMS samples were post-fixed with 3% paraformaldehyde + 0.1% glutaraldehyde in 1xPBS for 10 minutes followed by three rinses with 1xPBS.

### Preparation for STED imaging

For live cell STED imaging of U-2 OS Rtn4-Halo cells transiently overexpressing SNAP-Sec61β, cells were live labeled with 1 μM ATTO 590-chloroalkane and 1 μM of SNAP-Cell 647-SiR (New England Biolabs; S9102S) (final concentrations), diluted in warm media, for 1 hour at 37 °C. For the live STED imaging, U-2 OS and U-2 OS Rtn4-knockout cells both transiently overexpressing Halo-Sec61β were live labeled with 1 μM ATTO 590-chloroalkane, diluted in warm media, for 1 hour at 37 °C. All live cell STED samples were rinsed three times with warm media and allowed to recover for 1 hour at 37 °C following incubation with diluted dyes. The media was replaced with Live Cell Imaging Solution (Gibco; A14291DJ) supplemented with 15 mM glucose (Sigma-Aldrich; G5767). Cells were then imaged immediately while being maintained at 37 °C with 5% CO2.

### Preparation for single-molecule counting

U-2 OS Rtn4-mEGFP and U-2 OS Nup96-mEGFP cells were removed from flasks using trypsin, mixed together, and seeded into the lanes of a plasma cleaned ibidi slide. The next day, cells were fixed and labeled as described above for immunofluorescence. The samples were labeled with a custom-ordered anti-GFP nanobody conjugated to a single docking site for fluorogenic DNA PAINT imaging (Chung et al., 2022) (Massive Photonics). After washing out excess nanobody from the overnight labeling, samples were imaged immediately using 50 nM of Cy3B imager B in a PBS-based buffer (1x PBS, 500 mM NaCl, 1 mM Trolox, 20 mM Sodium Sulfite, pH 7.3-7.5). Trolox was stored in 1 M aliquots at -20 °C in DMSO.

### Preparation for 4Pi-SMS imaging

The following samples were labeled sequentially starting with the nanobody overnight at 4 °C, then the primary antibody overnight at 4 °C, and finally the secondary antibody for 1 hour at room temperature. U-2 OS transiently overexpressing mCherry-Sec61β were prepared for immunofluorescence as described above using a rabbit anti-Rtn4 primary (Abcam; ab47085) with a goat anti-rabbit secondary conjugated to Alexa Fluor 647 (Life Technologies; A21245) and a custom ordered FluoTag X4 anti-RFP nanobody conjugated to CF660C (NanoTag; N0404). U-2 OS transiently overexpressing GFP-KDEL or mEmerald-Sec61β were both prepared for immunofluorescence as described above using a rabbit anti-GFP primary (Invitrogen; A-11122) with a goat anti-Rabbit secondary conjugated to CF660C (Biotium; 20812) and a FluoTag X4 anti-GFP nanobody conjugated to Alexa Fluor 647 (NanoTag; N0304). To enable microscope alignment, 100 nm crimson Fluospheres (ThermoFisher; F8816) were added on top of all samples. Mounting the samples was performed as reported previously (Zhang et al., 2020).

### Microscopy

STED imaging was performed using a Leica SP8 STED 3x as described previously (Schroeder et al., 2019). A 100x 1.40 NA objective was used for all image acquisition using the Application Suite X Software (LAS X; Leica Microsystems). ATTO 590-chloroalkane and SNAP-Cell 647-SiR present in the same sample were imaged using excitation wavelengths of 592 nm (∼26 μW) and 650 nm (∼11 μW), respectively, and 775 nm (∼28 mW) depletion wavelength. HyD hybrid detectors with 0.3 – 6.0 ns gating were used to record fluorescence ranging from 600 – 630 nm for ATTO 590 and 650 – 750 nm for SNAP-Cell 647-SiR. Images were acquired sequentially between lines starting with SNAP-Cell 647-SiR using 8 line-average and 8000 Hz resonant scanning. One-color imaging of ATTO 590 in live cells was performed using 592 nm excitation at 24 μW, 775 nm depletion at 49 mW, 604 – 700 nm detection using a HyD hybrid detector with 0.3 – 6.0 ns gating, 8 line-average, and 8000 Hz scanning. Cells in all samples were maintained at 37 °C and 5% CO_2_. For figures, STED images were convolved with a Gaussian blur with a sigma of 1 pixel using ImageJ software. All analyses performed on STED images was done on the raw images.

4Pi-SMLM samples labeled with CF660C and Alexa Fluor 647 were imaged with a ratiometric approach using salvaged fluorescence on a 4Pi-SMLM microscope. The localization analysis and drift correction were executed in custom MATLAB scripts. All of this was performed as described previously (Zhang et al., 2020; Huang et al., 2016).

Single-molecule counting samples were imaged using Fluorogenic DNA PAINT on a Nikon Eclipse Ti2 microscope (Nikon Instruments) with an attached Andor Dragonfly 500 unit (Andor, Oxford instruments). Cy3B conjugated to fluorogenic imager probe B (Chung et al., 2022) at a working concentration of 50 nM was excited using a 561 nm laser with an 8.2 × 8.2 mm illumination aperture and a P2 power density filter resulting in a laser intensity of ∼1 mW/cm^2^. Fluorescence was filtered through a TR-DFLy-F600-050 emission filter before being collected using a 1.49 NA 60x oil TIRF objective and captured on Sona 4BV6X sCMOS camera (Andor Technologies) on 512 × 512 pixels with an effective pixel size of 108 nm at 20 Hz.

### ATTO 590 – Chloroalkane probe preparation

HaloTag chloralkane amine (2.2 mg, 9.9 μmol, 3.0 equiv) was prepared as previously described (Tyson et al., 2021) and added to a mixture of ATTO590-NHS (2.5 mg, 3.3 μmol, 1.0 eq), iPr2NEt (5.5 μL, 33 μmol, 10 eq) in DMSO. The reaction mixture was stirred for 3 h under the exclusion of light. Following completion of the reaction as tracked by LC-MS, the reaction mixture was purified directly via reverse-phase HPLC. The title compound was obtained as a solid after evaporation under reduced pressure of the collected HPLC fractions. HRMS(ESI): m/z calc. for C47H59ClN3O6+: 796.4087; found: 796.4073. UPLC:tR=1.88 min; gradient: 0 min: 5% B, 1 min:5% B, 1.6 min:95% B, 3.0 min: 95%B; C18 column).

### Skeleton generation and tubule major axis determination

To generate skeletons from a surface, we implemented a mean curvature skeletonization (Tagliasacchi et al., 2012) approach in PYMEVisualize (Marin et al., 2021). We used a weighting of 20 and 0.01 for the velocity and medial axis terms, respectively, to create the initial mesoskeleton. Then, the position of each point on the skeleton was changed to the average position of all points within 50 nm of that point. We applied an upper threshold of 200 to the density of skeleton points within 50 nm of any given skeleton point. Points exceeding this threshold were removed from the skeleton. This served to remove parts of the skeletons found within ER sheets, so analyses were restricted to tubules only. The major axis of the tubule at any given skeleton point was determined by calculating vectors pointing from each skeleton point to the nearest skeleton point that was a minimum of 10 nm away.

### Data analysis

#### Rtn4 pixel intensity

The average Rtn4 intensity per pixel along tubules and sheet edges was analyzed using ImageJ by creating 10-pixel wide regions of interest (ROI) along tubules. The Rtn4 pixel intensities were summed across the width of the line plots and those sums were averaged along the length of the ROI. The two populations of Rtn4 pixel values were tested for a statistically significant difference by computing the Wilcoxon rank-sum statistic.

#### Tubule diameter comparison

Tubule diameters were measured using 10-pixel wide line plots in Python Microscopy Environment (PYME) with the NEP fitting plug-in (Barentine et al., 2018). The “STEDTubule_SurfaceSNAP” model was used for all tubule diameter measurements. Pairs of tubule diameters populations were test for statistically significant differences by computing the Wilcoxon rank-sum statistic.

#### Mean curvature and Rtn4 density analysis

Three-dimensional surfaces of the ER were created in PYMEVisualize using the shrinkwrapping algorithm (Marin et al., in preparation) and the surfaces’ mean curvatures were calculated (Taubin) as implemented in PYMEVisualize (Marin et al., 2021). To determine the Rtn4 density near certain surface curvatures, the maximal principal curvatures along the surface were divided into 50 bins. The Rtn4 density was calculated as the number of Rtn4 localizations within 50 nm of a surface vertex (see **Fig. S2** for vertex description and visual) and were binned according to that vertex’s maximal principal curvature. “Rtn4 density/Frequency” was calculated by dividing the bin counts for Rtn4 density by the bin counts for the maximal principal curvature. Images were created using PYMEVisualize (Marin et al., 2021) and custom Python scripts.

#### φ calculation and simulated random data

The φ angles of Rtn4, Sec61β, and KDEL localizations in tubule cross-sections were calculated using a custom Python script. In PYMEVisualize, small regions of interest (ROIs) of ER tubules were created from larger datasets. The 3D surfaces for each ROI were generated based off the localizations from the nanobody labeling in each sample, and the skeletons were generated from those 3D-surfaces. Vectors pointing from the skeleton of the tubule to each localization from the secondary antibody labeling in each sample were calculated. The angles between these vectors and the z-axis (vector normal to the coverslip) were calculated using **Equation 1**. We used the skeletons from each dataset to simulate data randomly distributed around them in circular rings to show what the φ distribution would be for such data (**Fig. S3 C)**. The distributions of angles for simulated, Rtn4, Sec61β, and KDEL points were tested for significant differences by calculating the Wilcoxon rank-sum statistic.

#### Single-molecule counting analysis

Single-molecule counting datasets using fluorogenic DNA PAINT probes were localized in PYME. For each dataset, separate ROIs of Nup96 and Rtn4 were created. The Nup96 ROI was fed into Super-resolution Microscopy Platform (SMAP) (Thevathasan et al., 2019; Ries, 2020). Circular ROIs of individual nuclear pore complexes (NPC) were first automatically segmented in SMAP followed by manually removing out of focus ROIs. Built-in tools in SMAP were used to determine the effective labeling efficiency and the number of localizations per NPC. This information enables the calculation of the number of localizations per Nup96 protein as well as the number of localizations per labeled Nup96 protein. The Rtn4 ROI was segmented with DBSCAN, using a search radius of 15 nm and a minimum cluster size of 1, to identify unique clusters of Rtn4 localizations. Small clusters were later filtered out if they contained less localizations than the determined number of localizations per labeled Nup96 for that dataset, to ensure the clusters of points contained at least one protein. The number of Rtn4 proteins per cluster was calculated by dividing the number of Rtn4 localizations in each cluster by the number of localizations per Nup96.

#### ψ calculation

DBSCAN was used to segment 3D point-clouds of Rtn4 localizations into unique clusters using a search radius of 12 nm and a minimum clump size of 5. Clusters were analyzed using the ‘MeasureClusters3D’ module within PYMEVisualize. Specifically, principal component analysis was used to determine the principal axis of the clusters with the highest eigenvalue, which we refer to as the major axis. Vectors 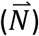 pointing from each cluster’s center to the closest point on the skeleton were calculated. Each cluster’s major axis was projected onto a plane orthogonal to its 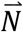. The ψ angles were calculated by using **Equation 1** with the projected major axes of the clusters and vectors orthogonal to the tubule axes and tangent to the projected plane, as input vectors. Only clusters containing more than 10 localizations and less than 50 localizations were used for this analysis. This removed background clusters and clusters that were too close together to segment, respectively. The relative distribution of different sized clusters is shown in **Fig. S6 A**.

#### Rtn4 3D cluster anisotropy analysis and simulated linear and circular clusters

Circular and linear clusters were simulated to compare to observed Rtn4 cluster. The simulated clusters possessed the same center points, radii of gyration (Rg), and localization counts as the observed Rtn4 clusters. To simulate circular clusters, each point in the cluster was assigned a uniform random angle from 0 to 2π and a uniform random radius (distance from the cluster center) varied as 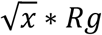 where x ranged from 0 to 1 and Rg was copied from the observed cluster each individual simulated cluster was based on. To simulate linear clusters, each point in the cluster was assigned the same angle, initially pulled from a uniform random distribution from 0 to 2π, and a uniform random radius from 0 to the radius of gyration. All points in both types of simulated clusters had some uncertainty incorporated into their positions. This was done by adding random numbers from a gaussian distribution (with μ = 0 and σ = 5.26; the average localization precision of the real data) to each component of their coordinates. The anisotropies of the simulated clusters and the observed Rtn4 clusters were calculated and compared. Clusters were filtered as 10 < localizations < 50, to avoid background clusters and clusters that were too close together to segment.

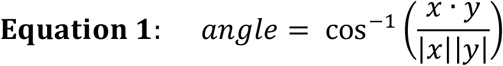

Where x and y are vectors and |x| and |y| are there magnitudes.

The Rg calculated from the observed clusters and used in the simulations are likely overestimates since they are effectively the true Rg convolved by the localization precision. Thus, our linear-shaped simulated clusters will appear slightly more anisotropic than they would if the true Rg were used. Indeed, if we multiply the calculated Rg by a factor of 0.925, the anisotropy distributions for the linear-shaped simulated data shift left and more closely align with the observed cluster data **(Fig. S8)**.

#### Simulating clusters with explicit inter-monomer distances

The frequency of different oligomer sizes was taken from the DNA-PAINT counting data **(Fig. 4)**. We determined a factor (3.23) that could transform the distribution of localizations per cluster of the 3D data to one that looked identical to that of the distribution of proteins per cluster from the counting data. This factor is roughly how many localizations we identify per protein. This was used as the λ for a Poisson distribution that was drawn from to determine the number of localizations per monomer in our simulated clusters. We added some localization precision to each simulated localization by adding some uncertainty drawn from a gaussian distribution with μ = 0 nm and σ = 5.26 nm, which is the average localization precision of our real data. The inter-monomer distances tested were based around the theoretical calculation of Rtn4b molecules possessing a diameter of ∼4.5 nm, assuming they are roughly spherical. The following website was used to calculate the volume of Rtn4b based off its sequence: http://biotools.nubic.northwestern.edu/proteincalc.html.

#### Statistics

All statistical tests were performed using the SciPy library of Python functions.

## Acknowledgements

This work was support by the National Institutes of Health (R01 GM118486, S10 OD020142, T32 GM007223, T32 EB019941) and the Wellcome Trust (203285/B/16/Z). The authors would like to thank Lena K. Schroeder for insightful discussions about endoplasmic reticulum structure as it pertains to Rtn4, Kenny K. H. Chung and Florian Schueder for the helpful discussions and expert advice regarding DNA-PAINT imaging, Jonas Ries for advice on best practices for single-molecule counting, Maohan Su for his assistance and expertise using the 4Pi-SMS microscope, and Felix Rivera-Molina (CINEMA Lab) for his assistance and expertise using the Andor Dragonfly microscope. We are grateful to Lena K. Schroeder and Shirin Bahmanyar for their thoughtful comments and feedback on the manuscript.

## Author contributions

L.A. Fuentes and J. Bewersdorf conceived the project. L.A. Fuentes was responsible for CRISPR cell line development, cell culture, sample preparation, image acquisition, and data simulations. Z. Marin developed the skeletonization method and implemented it in PYMEVisualize. L.A. Fuentes and Z. Marin designed and implemented the image analyses. J. Tyson synthesized the ATTO 590-Chloroalkane dye used for live cell STED imaging. D. Baddeley and J. Bewersdorf supervised image analysis, data simulations, and skeletonization development. L.A. Fuentes and J. Bewersdorf wrote the manuscript. All authors discussed the project and edited and reviewed the manuscript.

## Code availability

Python Microscopy Environment (PYME) is available at https://python-microscopy.org/. All custom scripts were created in jupyter-lab and are available at the GitHub repository: (https://github.com/lukasfue/Rtn4_paper_code).

## Abbreviations

ER: endoplasmic reticulum Rtn4: reticulon 4
STED: Stimulated Emission Depletion
SMLM: single-molecule localization microscopy Rg: Radius of gyration
DBSCAN: density-based spatial clustering of applications with noise

## Figures

**Figure S1.**
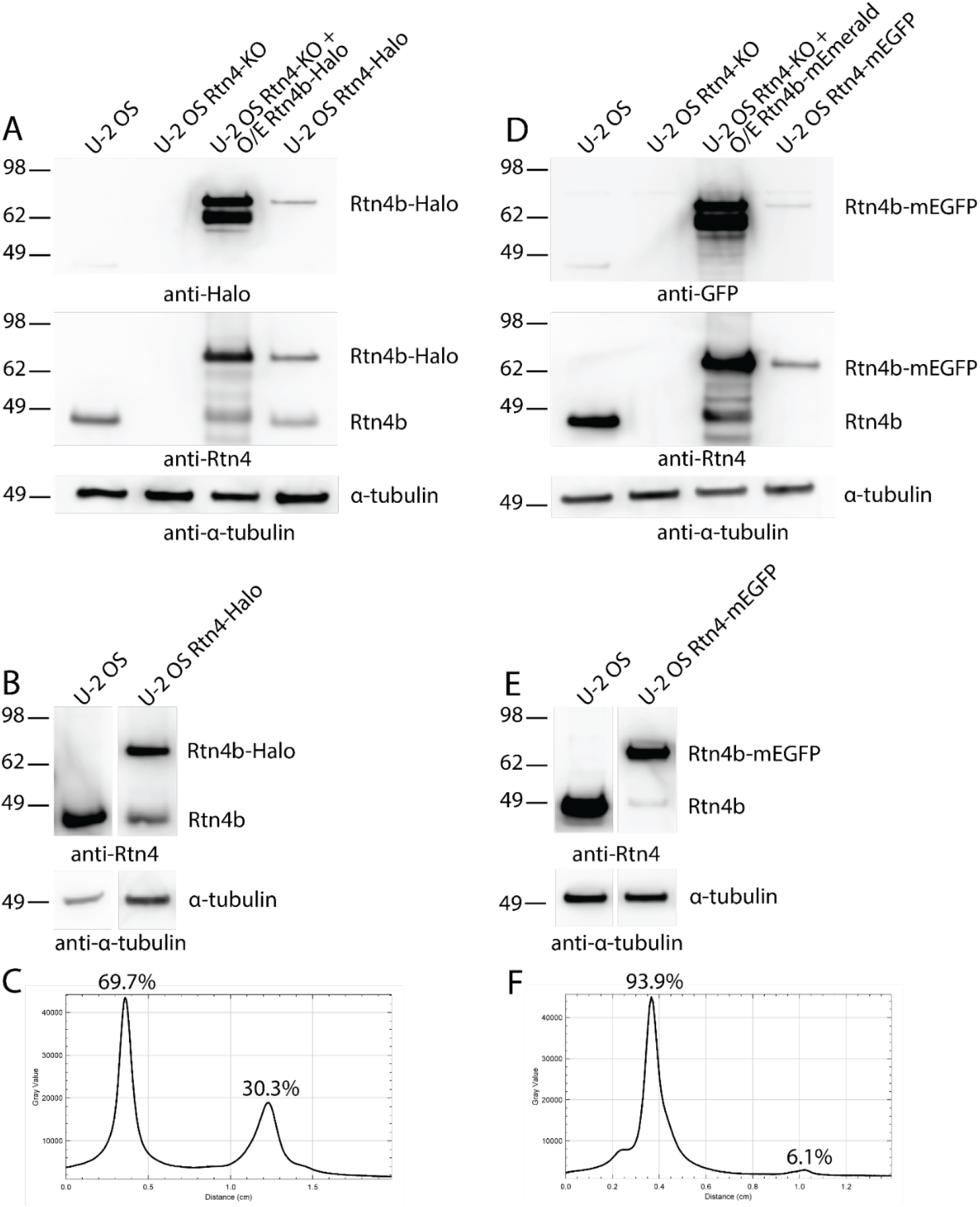
Western blot of CRISPR cell lines. (A) Probing U-2 OS Rtn4-Halo and control cell lines with anti-Halo, anti-Rtn4, and anti-α-tubulin antibodies. O/E stands for overexpressed. (B) Probing U-2 OS Rtn4-Halo cell line and WT U-2 OS cells only with anti-Rtn4 to avoid saturation from overexpressed proteins. (C) Line plot drawn through the U-2 OS Rtn4-Halo bands of tagged (top band/left peak) and untagged (bottom band/right peak) from blot in (B). (D) Probing U-2 OS Rtn4-mEGFP and control cell lines with anti-GFP, anti-Rtn4, and anti-α-tubulin antibodies. (E) Probing U-2 OS Rtn4-mEGFP cell line and WT U-2 OS cells only with anti-Rtn4 to avoid saturation from overexpressed proteins. (F) Line plot drawn through the U-2 OS Rtn4-mEGFP bands of tagged (top band/left peak) and untagged (bottom band/right peak) from blot in (E).

**Figure S2.**
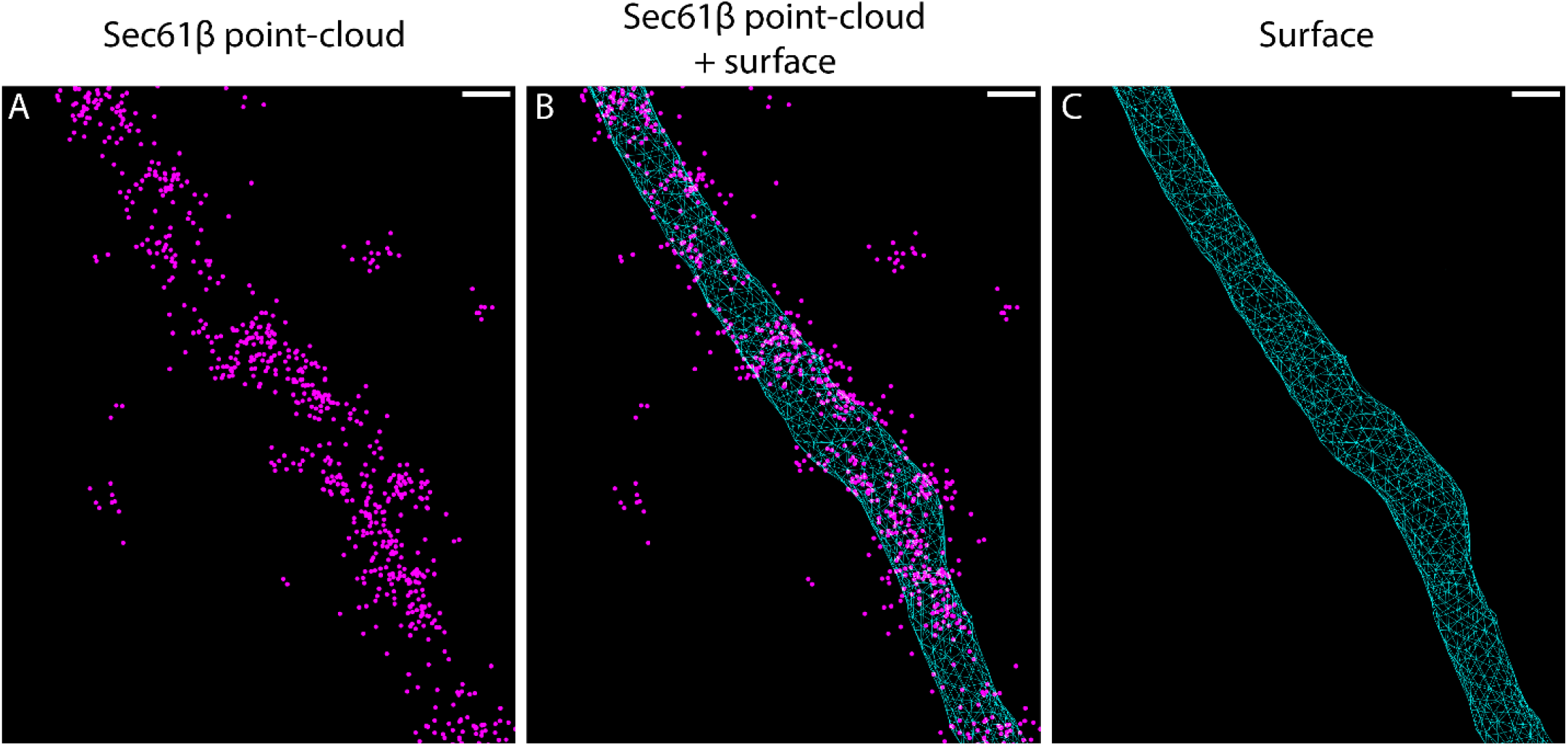
Point cloud surface fitting. (A) Example of a point-cloud of Sec61β. (B) Point-cloud in (A) and a surface generated from that point-cloud. (C) The surface alone. The surface is composed of triangles, which are joined by shared edges. We refer to the points where multiple edges meet as vertices and use their locations to calculate the local density of Rtn4. All scale bars represent 50 nm.

**Figure S3.**
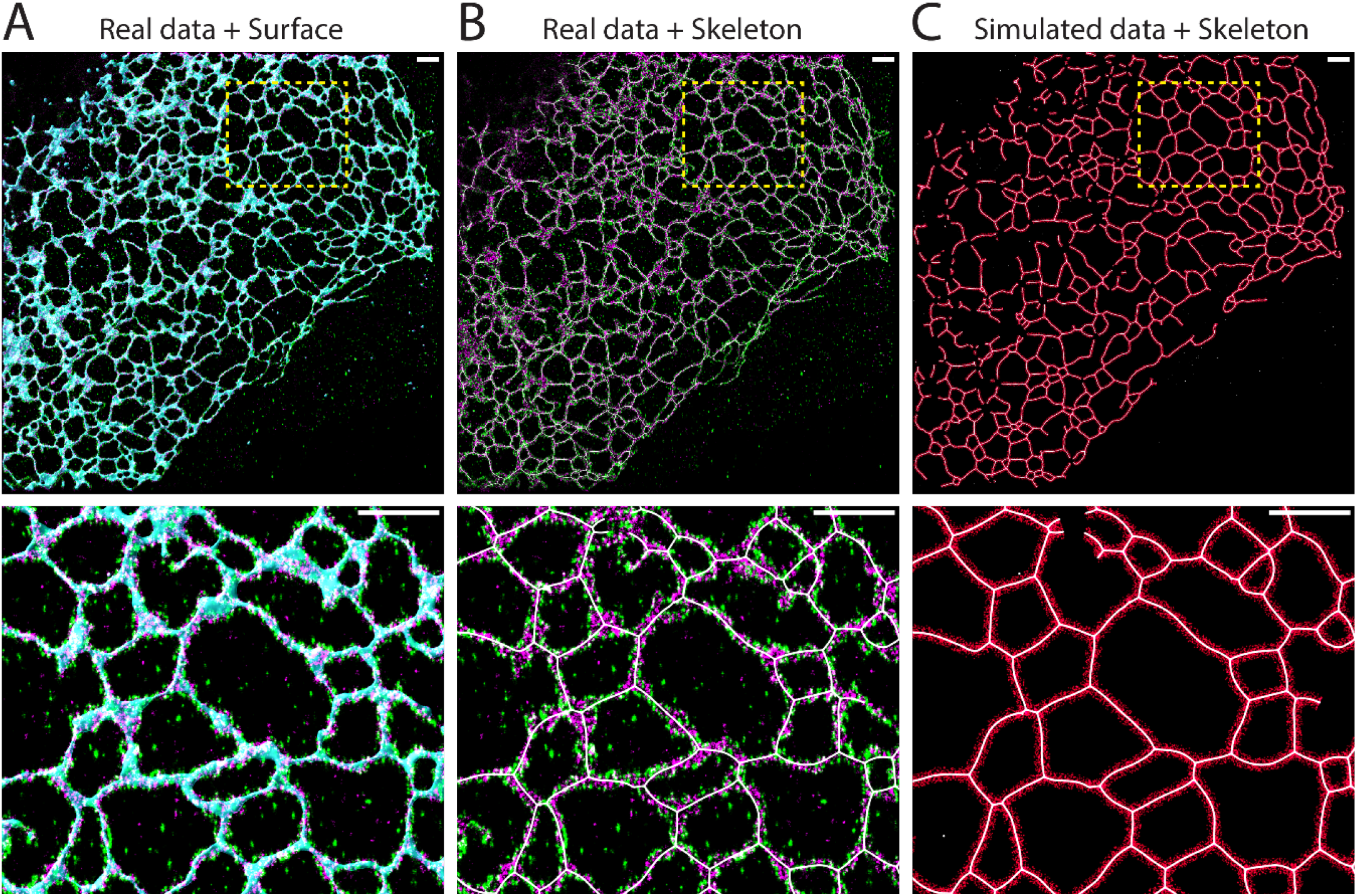
Skeletons and simulated data. (A) Two-color 3D SMLM data of Rtn4 (green), Sec61β (magenta), and a 3D surface based on the Sec61β point-cloud (cyan). (B) As in (A) but with the 3D surface removed and the skeleton (white) added. (C) Skeleton (white) is shown with data randomly simulated in circular rings (red) around the it. Scale bars represent 1 μm.

**Figure S4.**
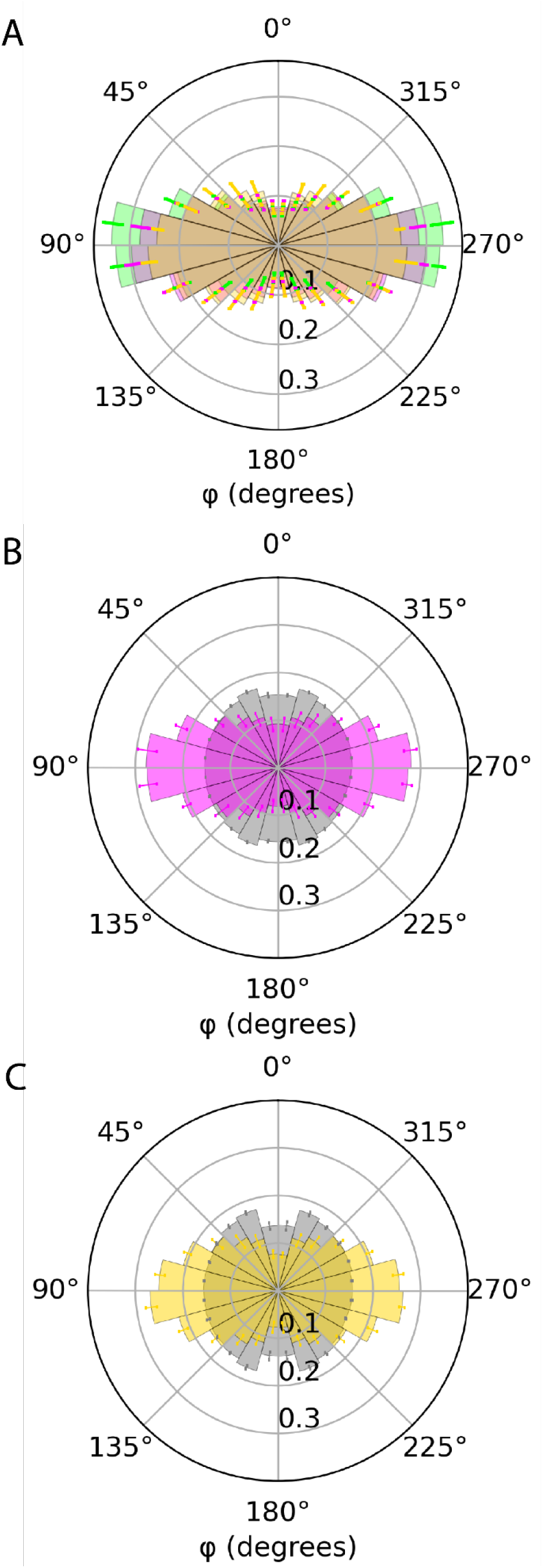
Control histograms of φ distributions. (A) The histograms of Rtn4 (green), Sec61β (magenta), and KDEL (yellow) together. The distributions were tested for significant differences using a rank sum test (Rtn4 vs. Sec61β: p<<<0.001; Rtn4 vs. KDEL: p<<<0.001; Sec61β vs. KDEL: p<<<0.001). (B) Polar histogram of φ from analysis using the nanobody localizations of Sec61β. (A) Polar histogram of φ from analysis using the nanobody localizations of KDEL.

**Figure S5.**
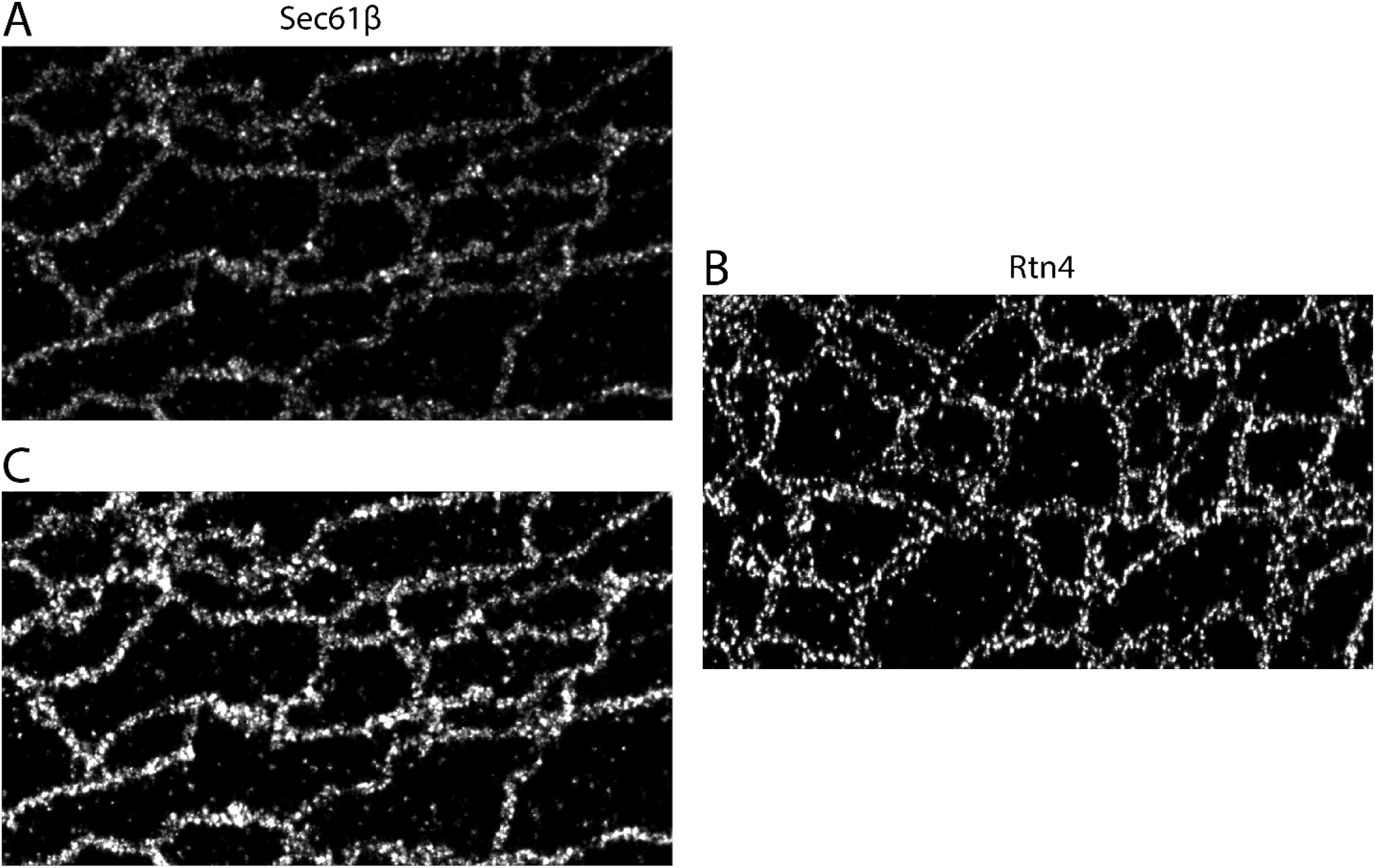
Comparison between Sec61β and Rtn4 localizations. (A, B) Sec61β and Rtn4 displayed with identical contrast settings. (C) The same image as (A) but displayed with adjusted contrast so that it can be more easily compared with (B).

**Figure S6.**
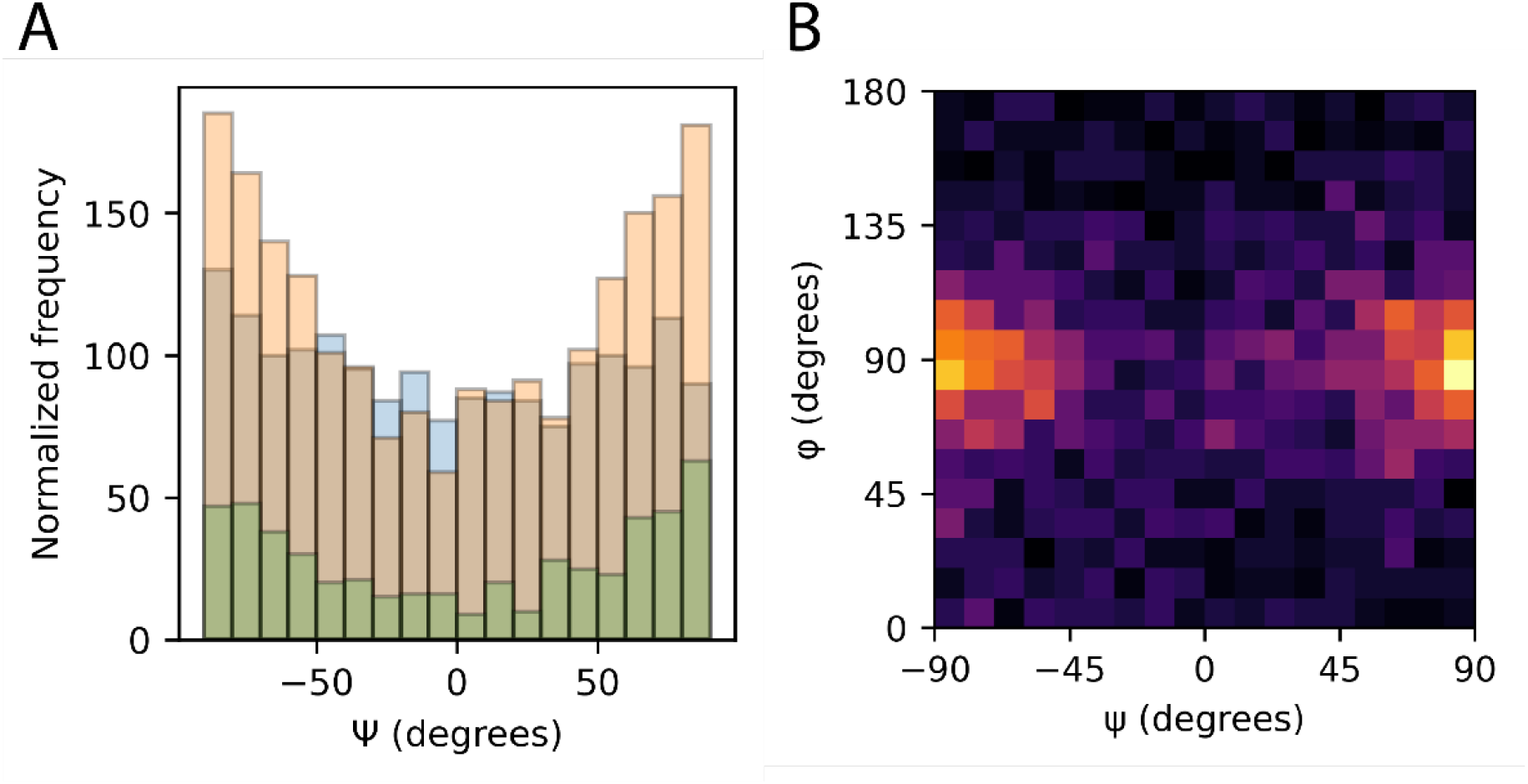
Distribution of different sized clusters’ ψ and how ψ and φ relate. (A) Histograms of ψ for small (blue; < 10 localizations), medium (orange; 10 < localizations < 50), and large (green; > 50 localizations) clusters. (B) 2D histogram of ψ vs. φ.

**Figure S7.**
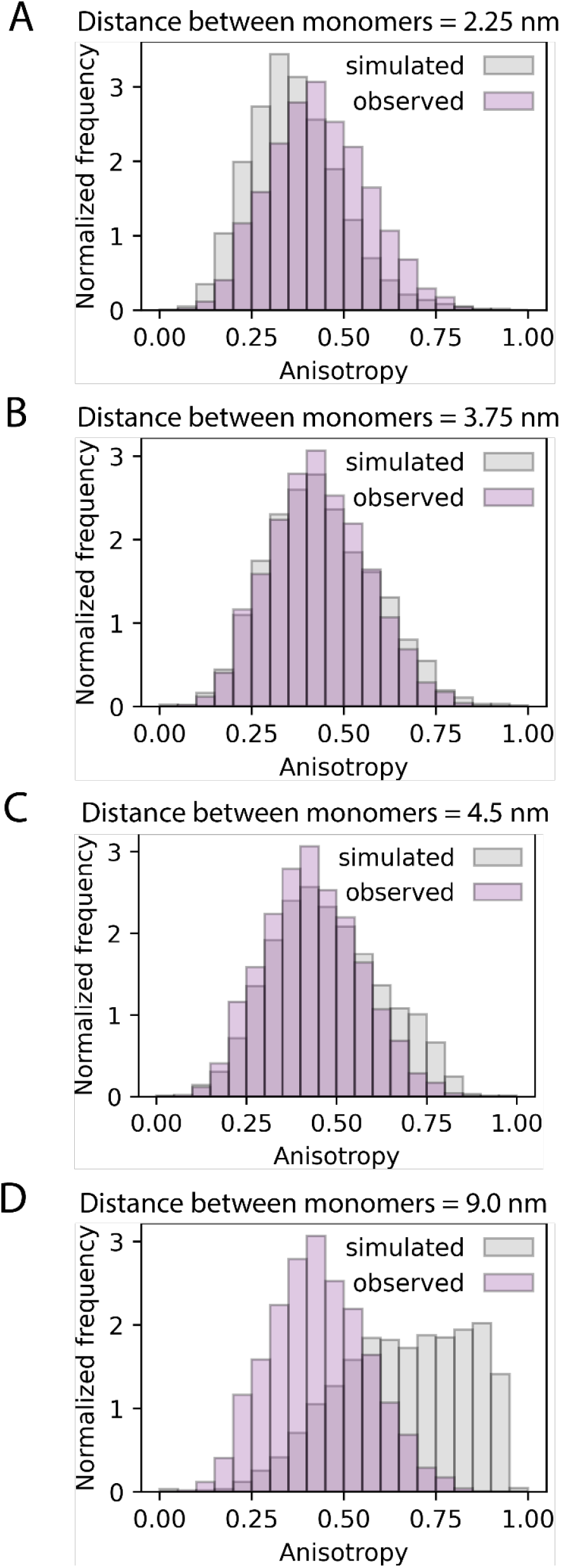
Anisotropy histograms of clusters simulated with explicit inter-monomer distances. (A-D) Histograms of observed cluster anisotropies (lavender) and clusters simulated with specific distances between the Rtn4 monomers (grey).

**Figure S8.**
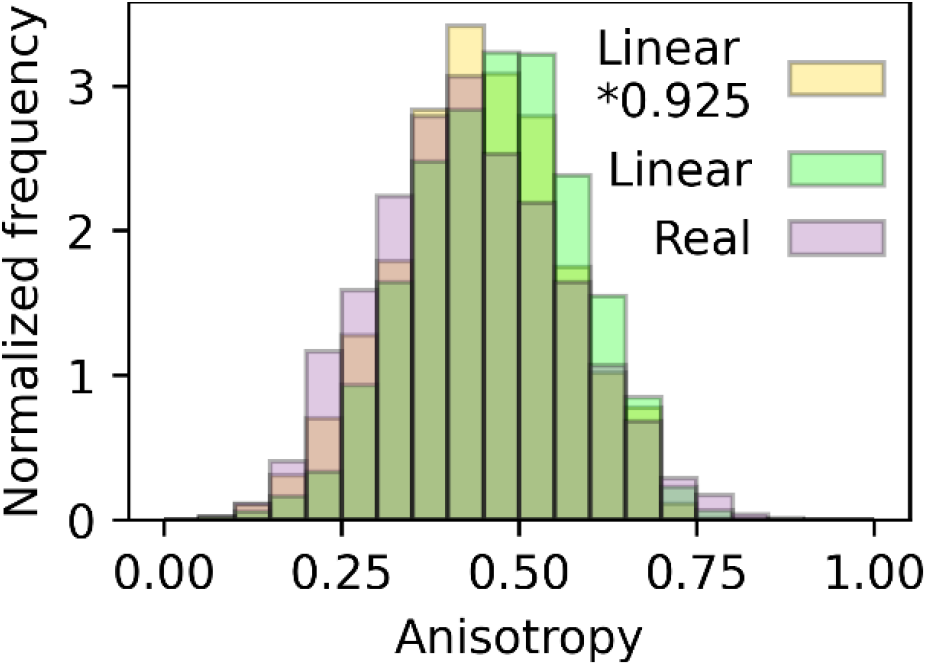
Corrected radius of gyration. Anisotropy distributions of observed clusters (lavender), linear-shaped clusters using the radius of gyration (Rg) of observed clusters (green), and linear-shaped clusters using Rg * 0.925 (yellow).

**Video 1** | **Rtn4 localizations on 3D surface**. Rtn4 localizations (green spheres) are shown on the 3D surface based on the mCherry-Sec61β point-cloud (shaded magenta surface).

**Video 2** | **Rtn4 clusters**. An example tubule showing the surface (cyan), skeleton (red line), the centers of each clusters (red circle), the major axis of each cluster (white line), and clusters of Rtn4 points (varying colors). Scale bar represents 50 nm.

**Video 3** | **Circular simulated clusters**. An example tubule showing the surface (cyan), skeleton (red), the centers of the clusters (red circle), and circular simulated clusters (yellow). Scale bar represents 50 nm.

**Video 4** | **Linear simulated clusters**. An example tubule showing the surface (cyan), skeleton (red), the centers of the clusters (red circle), and linear simulated clusters (green). Scale bar represents 50 nm.

**Video 5** | **Simulated cluster orientations**. An example tubule showing the surface (cyan), skeleton (red), the centers of the clusters (red circle), and the orientations of simulated clusters possessing all possible orientations tangential to the surface (white lines). Scale bar represents 50 nm.

